# Participant-derived cell line transcriptomic analyses and mouse studies reveal a role for ZNF335 in plasma cholesterol statin response

**DOI:** 10.1101/2023.06.14.544860

**Authors:** Elizabeth Theusch, Flora Y. Ting, Yuanyuan Qin, Kristen Stevens, Devesh Naidoo, Sarah M. King, Neil Yang, Joseph Orr, Brenda Y. Han, Jason G. Cyster, Yii-Der I. Chen, Jerome I. Rotter, Ronald M. Krauss, Marisa W. Medina

## Abstract

**Background:** Statins lower circulating low-density lipoprotein cholesterol (LDLC) levels and reduce cardiovascular disease risk. Though highly efficacious in general, there is considerable inter-individual variation in statin efficacy that remains largely unexplained.

**Methods:** To identify novel genes that may modulate statin-induced LDLC lowering, we used RNA-sequencing data from 426 control- and 2 μM simvastatin-treated lymphoblastoid cell lines (LCLs) derived from European and African American ancestry participants of the Cholesterol and Pharmacogenetics (CAP) 40 mg/day 6-week simvastatin clinical trial (ClinicalTrials.gov Identifier: NCT00451828). We correlated statin-induced changes in LCL gene expression with plasma LDLC statin response in the corresponding CAP participants. For the most correlated gene identified (*ZNF335*), we followed up *in vivo* by comparing plasma cholesterol levels, lipoprotein profiles, and lipid statin response between wild-type mice and carriers of a hypomorphic (partial loss of function) missense mutation in *Zfp335* (the mouse homolog of *ZNF335*).

**Results:** The statin-induced expression changes of 147 human LCL genes were significantly correlated to the plasma LDLC statin responses of the corresponding CAP participants *in vivo* (FDR=5%). The two genes with the strongest correlations were zinc finger protein 335 (*ZNF335* aka *NIF-1*, rho=0.237, FDR-adj p=0.0085) and CCR4-NOT transcription complex subunit 3 (*CNOT3*, rho=0.233, FDR-adj p=0.0085). Chow-fed mice carrying a hypomorphic missense (R1092W; aka bloto) mutation in *Zfp335* had significantly lower non-HDL cholesterol levels than wild type C57BL/6J mice in a sex combined model (p=0.04). Furthermore, male (but not female) mice carrying the *Zfp335*^*R1092W*^ allele had significantly lower total and HDL cholesterol levels than wild-type mice. In a separate experiment, wild-type mice fed a control diet for 4 weeks and a matched simvastatin diet for an additional 4 weeks had significant statin-induced reductions in non-HDLC (−43±18% and -23±19% for males and females, respectively). Wild-type male (but not female) mice experienced significant reductions in plasma LDL particle concentrations, while male mice carrying *Zfp335*^*R1092W*^ allele(s) exhibited a significantly blunted LDL statin response.

**Conclusions:** Our *in vitro* and *in vivo* studies identified *ZNF335* as a novel modulator of plasma cholesterol levels and statin response, suggesting that variation in ZNF335 activity could contribute to inter-individual differences in statin clinical efficacy.

## Background

Elevated plasma low-density lipoprotein cholesterol (LDLC) levels are a major risk factor for cardiovascular disease (CVD), the leading cause of death worldwide. Statins lower plasma LDLC levels and CVD risk by inhibiting cholesterol biosynthesis and stimulating receptor-mediated LDL uptake from the blood. Although statins are effective for primary and secondary prevention of atherosclerotic CVD [1], not everyone receives the same lipid-lowering benefit from statin therapy, and many individuals still experience CVD events while taking statins. Much of the inter-individual variation in statin efficacy remains unexplained to date.

Previous studies have identified some factors (e.g., age, smoking status, and race/ethnicity) associated with LDLC statin response [2]. In addition, genome-wide association studies (GWAS) have identified a handful of genetic loci associated with LDLC response [3-6], but together these only explain about 5% of LDLC response variation [5]. Though expanding sample sizes of participants with suitable pre- and on-drug phenotype data could bolster power for statin response GWAS, other approaches beyond GWAS could also be used to identify genetic factors that influence statin efficacy.

Statins elicit a strong transcriptional response via regulation of sterol response element binding factor 2 (SREBF2, aka SREBP2) [7] and other factors, so we and others have studied statin-induced gene expression changes to reveal novel genes involved in the modulation of clinical statin response [8-11]. To do this, we have used statin clinical trial participant-derived lymphoblastoid cell lines (LCLs) as an *in vitro* genetic model in which to study statin response transcriptomics. Here, we conducted a large-scale study correlating *in vitro* gene expression statin response with *in vivo* LDLC statin response to identify candidate statin efficacy modulators. We then validated a top candidate gene *in vivo* in a mouse model.

## Methods

### CAP study participants

This study used clinical and demographic information and cell lines derived from 426 participants in the Cholesterol and Pharmacogenetic (CAP) 40 mg/day six-week simvastatin clinical trial [2] (ClinicalTrials.gov Identifier: NCT00451828). Written informed consent was obtained from all participants, and the study was approved by the institutional review boards of the University of California, San Francisco and the University of California, Los Angeles. Plasma lipids were measured in two baseline (pre-statin) blood samples and two on-statin blood samples collected four and six weeks after statin initiation, and the two values were averaged in each case to obtain a single baseline and on-statin value per participant. LDLC values were calculated from total cholesterol, HDL-cholesterol, and triglyceride levels using the Friedewald formula [12]. Characteristics of this CAP subset are shown in **Table 1**. In addition, lipoprotein particle profiles were measured and analyzed in the second baseline sample and the 6-week on-statin sample from most CAP participants (n=803) using ion mobility (described below).

**Table 1.**
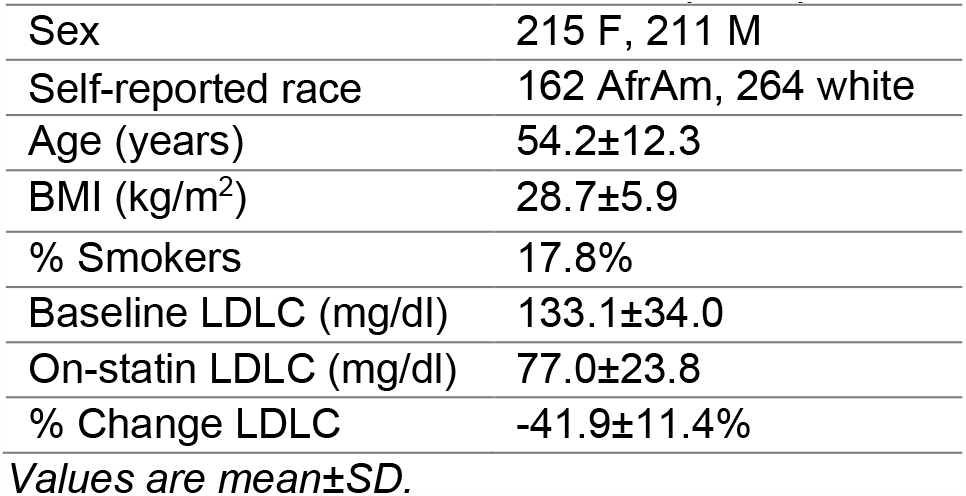
Characteristics of 426 CAP participants with LCL RNA-seq data.

### Lymphoblastoid cell line statin exposure

CAP lymphoblastoid cell lines (LCLs) were established by immortalizing blood-derived B lymphocytes through Epstein-Barr virus (EBV) transformation, cultured in RPMI 1640 with 10% FBS, and exposed to 2 μM activated simvastatin or sham control buffer for 24 hours in batches ≤12 cell lines, similar to previously described [13]. RNA was extracted from homogenized cellular lysates and RNA integrity was measured on an Agilent Bioanalyzer, as previously described [14].

### RNA-sequencing and analysis

PolyA-selected, strand-specific [15] paired-end RNA-seq libraries were prepared, and fragments were sequenced with 100-101 bp read lengths on Illumina machines at the University of Washington Northwest Genomics Center or the Baylor College of Medicine Human Genome Sequencing Center, as previously described [14, 16]. Sequences were aligned to the human (hg19) and EBV (NC_007605) genomes using Tophat v2.0.4 [17] with Ensembl v67 [18] and EBV [19] transcriptome annotations, and sequence fragments falling within annotated genes were counted using HTSeq [20] as previously described [14, 16]. DESeq2 was used to adjust for library size and to variance stabilize gene expression levels using a log_2_-based transformation [21], and statin-induced changes in gene expression levels were calculated by subtracting expression levels in control-treated LCLs from the statin-treated expression levels of the same cell line.

After adjusting CAP participants’ *in vivo* plasma LDLC statin responses [ln(on-statin LDLC)-ln(baseline LDLC)] for age, race/ethnicity, and smoking status, they were correlated to statin-induced changes in gene expression in the corresponding LCLs using Spearman’s rank-order correlation. P-values were adjusted for multiple testing using the Benjamini-Hochberg false-discovery rate (FDR) approach [22].

### Mouse studies

All mice in this study were housed under a 12-hour light/dark cycle and had free access to food and water except when fasted as noted. Mice were fed a general grain-based chow diet prior to study initiation (irradiated Teklad global 18% protein rodent diet from Envigo, product 2918). C57BL/6J mice carrying an N-ethyl-N-nitrosourea (ENU)-induced hypomorphic missense mutation (R1092W, aka “bloto”) in *Zfp335* were generated previously [23] and are available from JAX as stock #025853. For the first study, animals were fed the general chow diet until they were fasted for 4-6 hours and sacrificed at 13 weeks of age. Blood was collected via cardiac puncture, and plasma was isolated. Plasma lipid levels and lipoprotein particle concentrations were measured using Liasys and ion mobility, respectively, described below. Growth curve analyses were performed using weekly weights obtained after weaning, and growth curves of mice with different *Zfp335* genotypes were compared within sexes. Results were analyzed statistically and adjusted for multiple testing using the CGGC (Compare Groups of Growth Curves) method with 10,000 permutation tests [24, 25].

In our second study, mice were transitioned to the AIN-76A purified rodent diet (ResearchDiets product D10001i) at 13 weeks of age and weighed weekly for the duration of the study. After mice were fed the control purified diet for 4 weeks, they were transitioned to a matched diet with the addition of 1 gm simvastatin/kg diet (9.77% active; ResearchDiets product D14060903i) for 4 weeks. Plasma was obtained from mice at 15, 17, and 19 weeks of age using non-terminal retro-orbital bleeds after a 4-6 hour fast. Mice were sacrificed at 21 weeks of age after a 4-6 hour fast, and plasma was isolated from blood collected via cardiac puncture as in the first study.

Mouse studies were approved by the Children’s Hospital & Research Center Oakland and the University of California, San Francisco Institutional Animal Care and Use Committees.

### Plasma lipid analysis

Total cholesterol (TC) and high-density lipoprotein cholesterol (HDLC) were measured in human and mouse plasma samples on an AMS Liasys 330 Clinical Chemistry Analyzer, similar to previously described [26, 27]. Mouse non-HDL cholesterol (non-HDLC) levels were calculated by subtracting HDL from total cholesterol levels. In the chow-fed mouse experiment, HDLC was not successfully measured in samples from one male heterozygote mouse, and 1 female *Zfp335*^*R1092W*^ homozygote was a statistical (Tukey) outlier for HDLC (low) and non-HDLC (high) and was excluded from analyses. Plasma samples from 4 mice (1 male wild type, 1 male heterozygote, 1 female heterozygote, and 1 male *Zfp335*^*R1092W*^ homozygote) in the statin response experiment for which non-HDLC was calculated to be ≤0 and/or fasting total cholesterol levels were abnormally low (≤50 mg/dl) were Tukey outliers and were excluded from subsequent analyses. Additive relationships were tested for association using linear regression, and comparisons between wild-type and mutant homozygotes were conducted statistically using t-tests.

Ion mobility (IM) was used to measure human CAP participant and mouse plasma lipoprotein particle concentrations as a function of particle size, as previously described [28, 29]. Of the 25 male and 33 female mice for which IM data was generated before and on the statin diet, nine were statistically significant LDL particle concentration outliers (Tukey) and excluded from subsequent IM data analysis, leaving data from 22 male and 27 female mice. Specifically, five mice (1 female wild-type, 1 female heterozygote, 1 female Zfp335^R1092W^ homozygote, and 2 male wild-type) had abnormally high pre-statin LDL particle concentrations, three mice (1 female wild-type and 2 female heterozygotes) had abnormally high on-statin LDL particle concentrations, and one (male wild-type) mouse had abnormally low on-statin LDL particle concentrations. Data was binned into lipoprotein subfraction categories based on particle diameter definitions previously determined using human studies and comparisons with other methods [29]. Two-way ANOVA with Tukey’s multiple comparisons test was used to compare statin-induced changes in LDL subfraction particle concentrations across mouse genotype categories.

## Results

### Transcriptomic statin response in LCLs

Statin induced a robust transcriptional response in the 426 CAP LCLs, in agreement with our previous studies [9, 14] (**Figure 1, Table S1**). Of the 14,028 well-expressed human and EBV genes tested, the expression level of 9,092 genes (65%) changed significantly with statin treatment (paired t-test, FDR-adjusted p<0.0001, 4,671 up-regulated and 4,421 downregulated), though the degree of change was modest for most genes. As expected, many of the most robustly up-regulated genes were known SREBF2 target genes.

**Figure 1.**
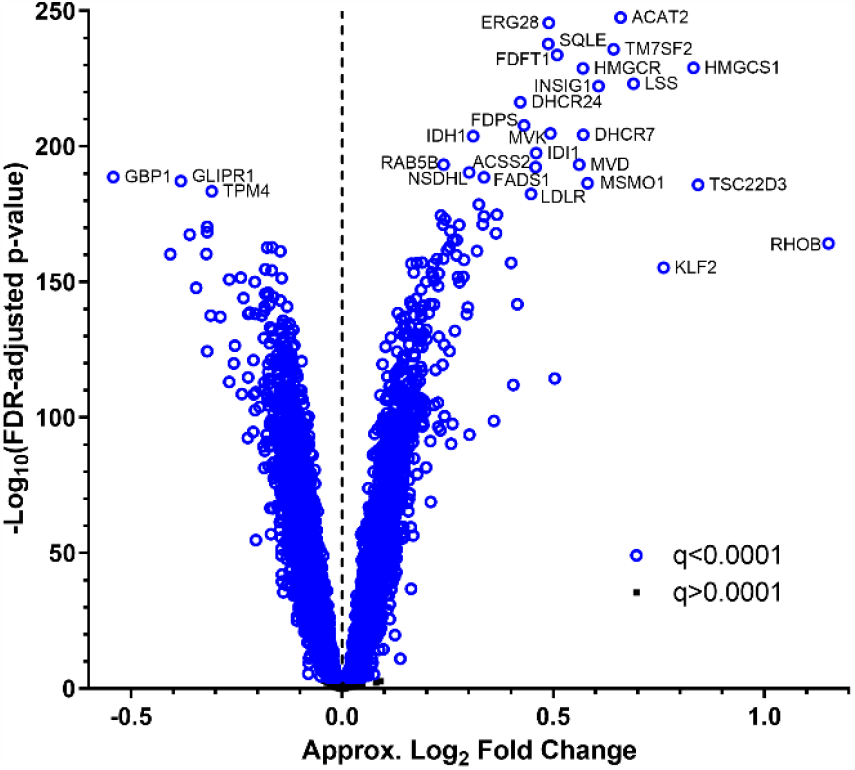
Volcano plot illustrating gene expression statin response in 426 CAP LCLs. Approximate log_2_ fold change is 2^(variance stabilized statin gene expression level – variance stabilized control gene expression level) calculated for each cell line and averaged across cell lines.

### Correlation of *in vitro* gene expression statin response with *in vivo* LDLC statin response

To identify genes that may modulate plasma LDLC statin response, we correlated transcriptome-wide statin-induced changes in LCL gene expression *in vitro* with plasma LDLC statin responses of the corresponding CAP clinical trial participants from whom the LCLs were derived. At a false discovery rate of 5%, 147 of 13,955 human genes tested had statin-induced expression changes that were significantly correlated with *in vivo* LDLC statin response (**Table S2**). The top 10 genes, which all had an FDR-adjusted p<0.03, are shown in **Table 2**.

**Table 2.** Genes with LCL expression changes most correlated to plasma LDLC statin response.

| Gene symbol | Gene name | Chr | Rho | P | Q |
| --- | --- | --- | --- | --- | --- |
| **ZNF335** | zinc finger protein 335 | 20 | 0.237 | 7.8E-07 | 0.0085 |
| **CNOT3** | CCR4-NOT transcription complex subunit 3 | 19 | 0.233 | 1.2E-06 | 0.0085 |
| **PML** | PML nuclear body scaffold | 15 | 0.218 | 5.4E-06 | 0.0246 |
| **SYVN1** | synoviolin 1 | 11 | 0.216 | 7.0E-06 | 0.0246 |
| **ANKRD46** | ankyrin repeat domain 46 | 8 | -0.208 | 1.6E-05 | 0.0293 |
| **EIF4E** | eukaryotic translation initiation factor 4E | 4 | -0.206 | 1.9E-05 | 0.0293 |
| **MED15** | mediator complex subunit 15 | 22 | 0.206 | 1.9E-05 | 0.0293 |
| **TCF3** | transcription factor 3 | 19 | 0.205 | 2.1E-05 | 0.0293 |
| **OTUD6B** | OTU deubiquitinase 6B | 8 | -0.205 | 2.1E-05 | 0.0293 |
| **AASDH** | aminoadipate-semialdehyde dehydrogenase | 4 | -0.205 | 2.1E-05 | 0.0293 |

*Rho* is the Spearman correlation coefficient and *Q* is the FDR-adjusted *P*-value.

The top two genes, zinc finger protein 335 (*ZNF335*) and CCR4-NOT transcription complex subunit 3 (*CNOT3*), appeared to be most worthwhile for follow-up due to the significance of their correlation relationships relative to those of other genes after multiple testing adjustment. ZNF335 (aka NIF-1) has been shown to physically interact with two other members of the CCR4-NOT complex, CNOT6 and CNOT9 [30], suggesting that ZNF335 and CNOT3 may be functionally related. *ZNF335* and *CNOT3* also exhibited similar correlation directionality (**Figure 2**). For both genes, statin decreased expression in most LCLs. In addition, the LCLs that exhibited greater statin-induced reductions in gene expression tended to be derived from participants who experienced larger statin-induced reductions in plasma LDLC in the CAP clinical trial (**Figures 2A-B**). Notably, the statin-induced changes in *CNOT3* and *ZNF335* were themselves highly correlated (rho=0.625, p=1.4×10^−47^, **Figure 2C**). Other genes in the CCR4-NOT complex did not have gene expression statin responses that were significantly correlated to plasma LDLC response after adjustment for multiple testing, and their non-significant correlations with plasma LDLC response were negative, displaying opposite directionality to the positive *CNOT3* and ZNF335 correlations (**Table S2**).

**Figure 2.**
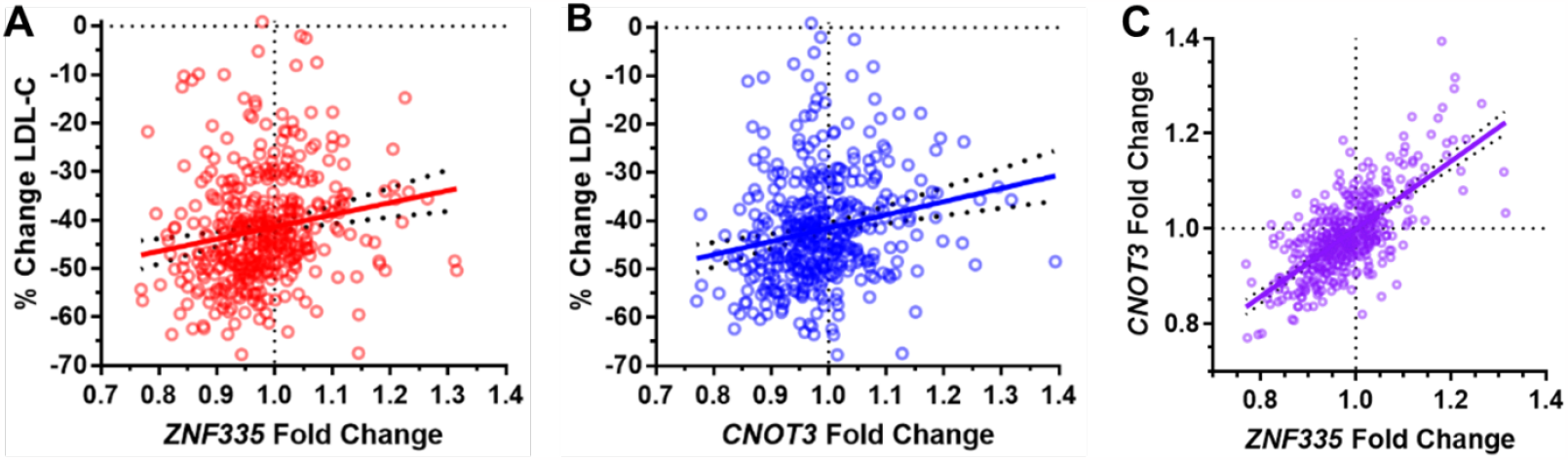
Correlation of *ZNF335* and *CNOT3* LCL gene expression statin responses with *in vivo* plasma LDLC statin responses of the corresponding donors (A-B) and with each other (C).

### Plasma cholesterol levels in *Zfp335* hypomorphic mice

In mice, knocking out *Cnot3* or *Zfp335* (the murine *ZNF335* homolog) results in embryonic lethality [31, 32]. Previous studies have shown that *Cnot3*^+/-^ (haplodeficient) mice have altered lipid and glucose metabolism [33], and liver-specific *Cnot3* knockout (Cnot3LKO) mice have altered lipid metabolism, among other deficits [34]. While ZNF335 has been reported to play a role in neurogenesis [32], to date the effects of *Zfp335* disruption on lipid and cholesterol metabolism have not previously been studied. We chose to investigate plasma cholesterol levels and statin response in mice carrying an ENU-induced hypomorphic missense mutation in *Zfp335* (R1092W aka bloto) that was previously studied in the context of T lymphocyte maturation [23].

We first compared fasting plasma cholesterol levels of 12-week-old *Zfp335*^*R1092W/R1092W*^, *Zfp335*^*R1092W/+*^, and wild type mouse littermates fed a grain-based chow diet. In male mice, increasing copies of *Zfp335*^*R1092W*^ mutations significantly reduced total cholesterol (TC) and HDL cholesterol (HDLC) levels (**Figure 3**, additive p=0.01 and p=0.0002, respectively). As expected based on previous studies, female mice had substantially lower HDL (and total) cholesterol levels than male mice across all *Zfp335* genotype categories. However, associations of *Zfp335* genotype with plasma TC or HDLC were not observed in female mice, with sex\**Zfp335* genotype interactions of p=0.15 and p=0.015, respectively. Since we observed a significant sex interaction for HDLC levels in the mice, we went back to our CAP participant data and observed a modest association of *ZNF335* levels in control-treated LCLs with CAP baseline HDLC in men (p=0.048) but no association in women, with a weak sex\**ZNF335* interaction (p=0.09, **Figure S1**). The directionality of the correlation in men seems consistent with the male mice as well, since lower human LCL gene expression and more copies of the mouse hypomorphic Zfp335 allele would both be presumed to have less protein activity.

**Figure 3.**
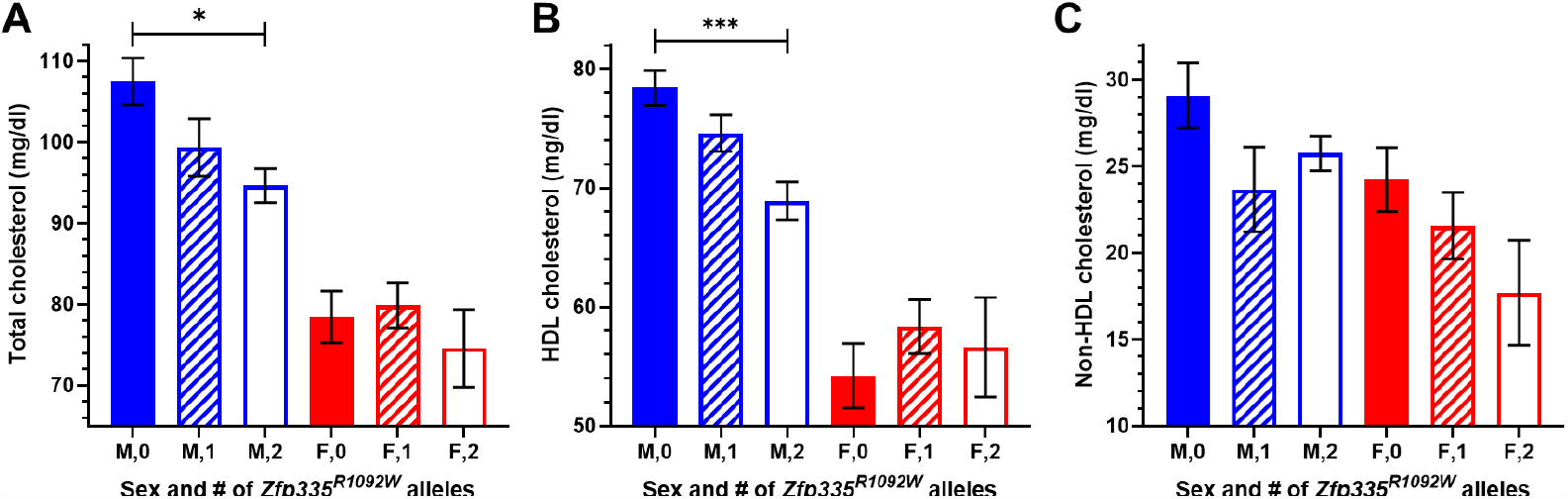
12-week old chow-fed mouse plasma (A) total, (B) HDL, and (C) non-HDL cholesterol levels split by sex and *Zfp335* genotype. Male sample sizes were N=12, 15 (16 for TC), and 10, and female sample sizes were N=17, 15, and 5 (6 for TC) for wild type, heterozygotes, and homozygous Zfp335R1092W, respectively. Values are mean ± SEM. *p<0.05 ***p<0.001

In contrast to HDLC, there were not significant sex differences in the relationship between mouse *Zfp335* genotype and non-HDL cholesterol (TC - HDLC). *Zfp335* genotype was significantly correlated with non-HDLC in a sex-combined model (additive p=0.04) with sex included as a covariate. Though not statistically significant, male mice with one or two copies of the *Zfp335*^*R1092W*^ mutation had lower mean levels of non-HDLC compared to wild type littermates. In female mice, the strongest relationship of genotype with cholesterol levels was for non-HDLC, with a trend for reduced levels with increasing numbers of *Zfp335*^*R1092W*^ mutations (additive p=0.08).

### Weight and growth curves

Since we observed that *Zfp335*^*R1092W/R1092W*^ mice appeared smaller on average than their littermates and this phenotype had not been documented for this mutant mouse strain before, we measured their growth curve from 4-11 weeks of age on a grain-based chow diet (**Figure S2**). Both male (adj. p=0.0036) and female (adj. p=0.036) *Zfp335*^*R1092W/R1092W*^ mice had significantly reduced growth curves compared to their wild-type littermates. For males, (which had a larger sample size than females) *Zfp335*^*R1092W/+*^ animals had growth curves that were intermediate between *Zfp335*^*R1092W/R1092W*^ and *Zfp335*^*+/+*^ males (heterozygote vs. *Zfp335*^*R1092W/R1092W*^ adj. p=0.0021, heterozygote vs. wild-type adj. p=0.0498), with females trending similarly (adj. p=0.098 for both comparisons).

### LDL statin response in *Zfp335* hypomorphic mice

We next wanted to determine if statin efficacy differed between wild-type mice and mice carrying *Zfp335*^*R1092W*^ mutations. To do this, we first fed mice a defined “control” diet (AIN-76A) for 4 weeks, starting at 13 weeks of age. We then transitioned mice to a matched diet supplemented with simvastatin for 4 weeks. We first measured statin-induced changes in non-HDLC as a crude proxy for mouse LDLC statin response, though non-HDLC also includes cholesterol found in other lipoprotein particles. Pre-statin non-HDLC levels (the average of the 15- and 17-week samples; 2 and 4 weeks on AIN-76A diet) were associated with *Zfp335* genotype in male animals alone (additive p=0.022, **Figure 4A**) and in a sex-combined model with sex included as a covariate (additive p=0.029), with a similar but non-significant trend in female animals alone (**Figure 4B**).

**Figure 4.**
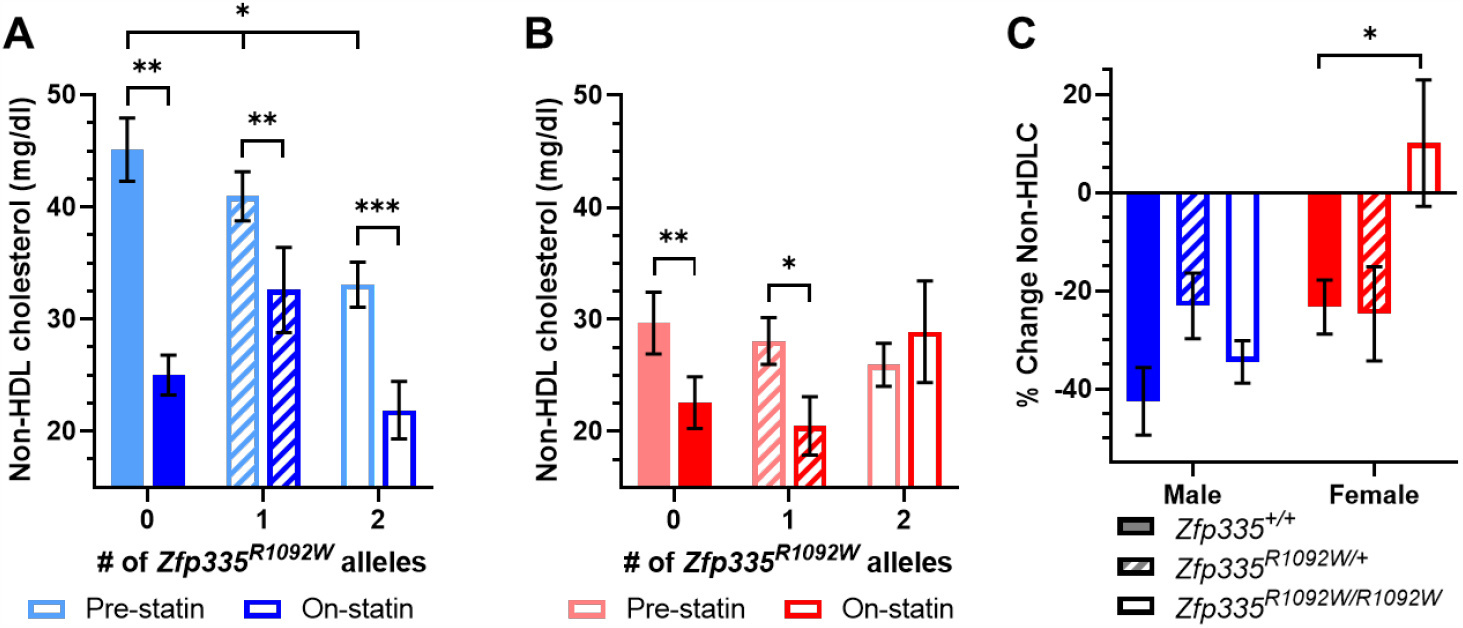
Plasma non-HDL cholesterol levels in (A) male and (B) female mice before and during statin diet feeding and (C) statin-induced changes in non-HDL cholesterol split by sex and *Zfp335* genotype. Male sample sizes were N=7, 15, and 4 and female sample sizes were N=12, 11, and 5 for wild type, heterozygotes, and homozygous Zfp335R1092W, respectively. Values are mean ± SEM. *p<0.05.

Non-HDLC levels were reduced after 4 weeks of statin treatment in wild-type mice of both sexes, with wild-type male mice experiencing greater (inter-sex comparison p<0.05) statin-induced reductions in non-HDLC (**Figures 4A and 4C**, -43±18%, paired t-test p=0.0016, N=7) than wild-type females (**Figures 4B-C**, -23±19%, paired t-test p=0.0020, N=12). Female *Zfp335*^*R1092W/R1092W*^ mice (N=5) had a significantly blunted non-HDL cholesterol statin response compared to their wild-type littermates (**Figure 4C**, p=0.02, t-test), but this trend was not as strong in male animals.

### Lipoprotein subfraction statin response

Non-HDL cholesterol is a broad category that includes cholesterol content from lipoprotein particles of various sizes and types. Since LDLC response was the phenotype for which we saw a correlation with *ZNF335* gene expression statin response in humans, we next performed a more detailed analysis of lipoprotein particles in the LDL size range. We therefore compared the ion mobility-based lipoprotein profiles from wild-type mice before and after they were fed 4 weeks of statin diet (**Figures 5 and S3**). As has been well documented previously, most mouse lipoproteins were in the HDL size range, with much lower concentrations in the size ranges spanning LDL and intermediate density lipoproteins (IDL). Since the lipoprotein subfraction size categorization we used is based on human data, it may not always accurately reflect the type(s) of particles in the corresponding size intervals in mice. In mice, particles in the human very small LDL (designated LDL IV) particle size range comprise a heterogeneous mixture of species overlapping very large HDL and containing APOE-only lipoproteins [27], so we focused our analyses on small (LDL IIIb [20.49-20.82 nm] and LDL IIIa [20.82-21.41 nm]), medium (LDL IIb [21.41-22 nm] and LDL IIa [22-22.46 nm]), and large (LDL I [22.46-23.33 nm]) LDL size ranges. Wild-type males were the only mice that exhibited robust statin-induced decreases in these LDL particles (**Figures 5A, 6, S3, and S4**). For comparison, we also included pre-statin and on-statin human lipoprotein profiles from CAP clinical trial participants (**Figure S5**). Similar to male mice, men experienced a significant reduction in lipoprotein particle concentrations in the medium and large LDL to small IDL size range after statin treatment, but unlike the sex difference in statin response in the mice, there were similar reductions in levels of these particles in women.

**Figure 5.**
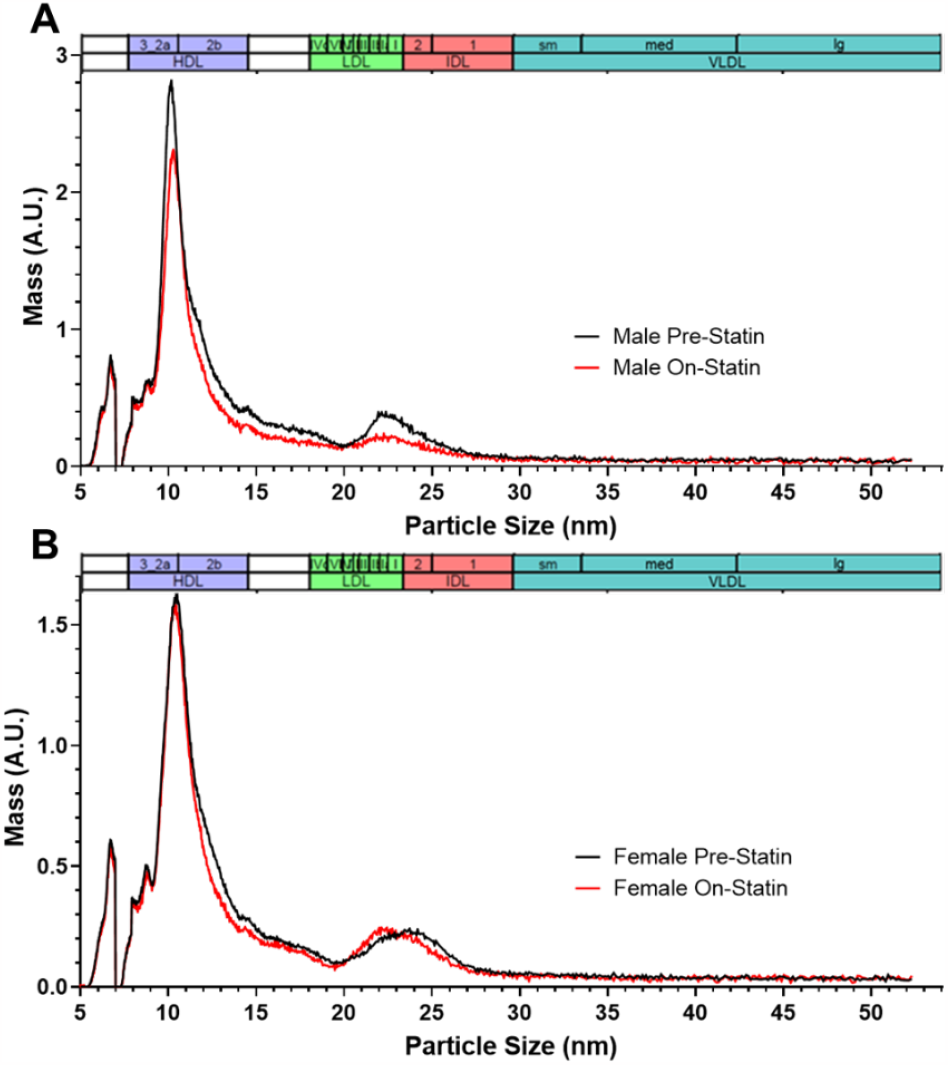
Statin-induced changes in wild-type A) N=5 male and B) N=11 female mouse lipoprotein profiles measured by ion mobility. Mouse profiles were measured before and after 4 weeks of simvastatin-containing diet. The size intervals designating the major lipoprotein subclasses are based on those defined in humans (29).

Male mice carrying one or two copies of the *Zfp335*^*R1092W*^ mutation exhibited significantly blunted LDL statin responses compared to wild-type (**Figure 6**, p=0.0011 and p=0.0010, respectively). Female mice of any genotype experienced minimal changes in LDL particle concentrations with statin treatment (**Figures 5B, S3, and S4B**); the statin-induced reductions in non-HDLC in female wild-type and heterozygous mice (**Figure 4C**) seemed largely driven by particles in the IDL size range instead (**Figures 5B, S3, and S4B**). We also went back to investigate whether there was a significant sex interaction of the original association of LCL *ZNF335* statin response with plasma LDLC statin response in CAP. This association was also stronger in men than women, though the interaction of sex with gene expression change was not significant (p=0.28, **Figure S6**).

**Figure 6.**
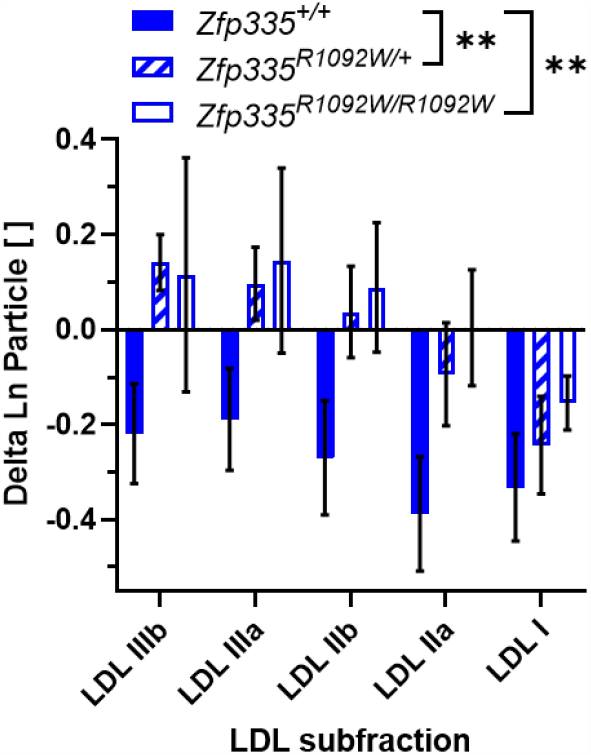
Statin-induced changes in male mouse LDL particle concentrations as measured by ion mobility. Sample sizes were N=5, 12, and 5 for wild type, heterozygotes, and homozygous Zfp335^R1092W^, respectively. Values are mean ± SEM. Genotypes were compared using two-way ANOVA with Tukey’s multiple comparison test. **p<0.01.

## Discussion

In this study, we show evidence that zinc finger protein 335 (*ZNF335*/*Zfp335*) is a novel modulator of LDLC statin response. In our *in vitro* experiments, we identify 147 human genes whose *in vitro* statin responses correlate significantly with the *in vivo* plasma LDLC statin responses of the corresponding donors, with *ZNF335* and *CNOT3* topping the list. In our subsequent *in vivo* experiments in mice using a *Zfp335* hypomorphic missense (*Zfp335*^*R1092W*^) genetic model, we show for the first time that reduced ZFP335 function leads to reduced plasma cholesterol levels and a blunted LDL response to statin, with significant sex differences.

ZNF335 resides in the nucleus and regulates transcription, including enhancing transcriptional activation by nuclear hormone receptors [35, 36]. It can activate transcription by recruiting histone methyltransferase complexes to the promoters of genes [32]. ZNF335 is widely expressed across human tissues [37] and has diverse functions, including roles in neuronal development [32] and T-cell maturation [23, 38, 39]. Here, we demonstrate for the first time that ZNF335 influences plasma cholesterol levels and statin response. Further investigation is needed to determine which, if any, genes affecting cholesterol metabolism may be regulated transcriptionally by *ZNF335* and to determine the mechanism by which *ZNF335* influences plasma cholesterol phenotypes.

ZNF335 physically interacts with members of the CCR4-NOT (carbon catabolite repression 4– negative on TATA-less) complex, including CNOT6 (aka CCR4) and CNOT9 (aka RCD1) [30]. The CCR4-NOT complex can regulate gene expression via several mechanisms, including transcription and mRNA deadenylation [40]. Though a physical interaction has not been reported between ZNF335 and the CCR4-NOT complex regulatory subunit CNOT3, here we observe strong correlations between their statin-induced gene expression changes, indicating they may be co-regulated by some of the same statin-responsive factors. *ZNF335* and *CNOT3* expression changes in LCLs also show similar strength and directionality of correlation with plasma LDLC statin responses of the corresponding statin clinical trial participants, so they may be functionally related, and it is plausible that they could be acting as components of the same complex. CNOT3 has been previously reported to have roles in regulation of lipid and glucose metabolism and to be differentially regulated in fasted and fed states based on studies in CNOT3 haplodeficient mice [33]. A functional link between ZNF335 and CNOT3 could be further explored in future studies, as could effects of conditional knockdowns of both genes on lipids and statin response.

As with *Cnot3* [31], knockout of *Zfp335* in mice is embryonic lethal [32]. Thus, to study the potential role of *Zfp335* in cholesterol metabolism and statin response, here we use mice carrying a hypomorphic mutation (R1092W aka bloto) in *Zfp335*. Previous studies using *Zfp335*^*R1092W*^ mice have shown that the R1092W mutation in the twelfth zinc finger of ZFP335 impairs its ability to bind a subset of its DNA targets [23] by modifying one of its two known DNA binding domains [41]. Previous studies of this genetic mouse model have focused on the role of ZFP335 in T-cell development, but here we also show that the mutation has an impact on body weight, blood cholesterol levels, and blood cholesterol statin response. It is interesting to note that for some of these phenotypes (e.g., body weight and HDL-cholesterol levels in male mice), heterozygotes display intermediate phenotypes typical of an additive relationship, while for other phenotypes, heterozygotes are phenotypically more similar to wild-type animals (e.g., non-HDL cholesterol statin response in female animals) or to *Zfp335*^*R1092W/R1092W*^ homozygous mutant animals (e.g., LDL particle statin response in male animals). Perhaps ZFP335 acts as a component of distinct complexes to elicit some of these phenotypes, which could explain why the same genetic model is not common to them all, or perhaps the fact that the mutation is only a partial loss of function mutation plays a role.

We observe striking sex differences when studying the impact of the hypomorphic missense *Zfp335* mutation on mouse HDL cholesterol levels and LDL statin response. Similar, though less significant, sex differences are also observed for the human LCL *ZNF335* statin response correlation with plasma LDLC statin response and the correlation of LCL *ZNF335* levels with plasma HDL-cholesterol levels, showing inter-species replication despite known differences in lipoprotein metabolism and profiles between the species. The differences in the impact of the *Zfp335* mutation between sexes could be related to sex-specific differences in the hormonal milieu, given ZNF335’s role in nuclear hormone receptor signaling, but the hormone(s) involved and which of their downstream targets regulate cholesterol levels and statin response remain to be elucidated.

Sex differences are also observed in the lipoprotein profiles and lipoprotein response to statin in wild-type mice. Wild-type female mice have less plasma HDL cholesterol and total cholesterol than males and exhibit a blunted non-HDL cholesterol statin response compared to males. In this study, though male mice experience significant reductions in LDL with statin treatment, the statin-induced reduction in non-HDL cholesterol in female mice appears to be driven mainly by larger particles (including IDL) rather than LDL. In contrast, in humans, women experience LDL reductions quite similar to men.

This study illustrates the utility of clinical trial participant-derived cell lines in identifying novel genetic factors involved in drug response phenotypes. The correlation of *in vitro* drug-induced transcriptomic changes in cell lines to *in vivo* drug response phenotypes from the corresponding participants during the clinical trial can be a powerful approach, even in the transformed cell lines used here. The emergence of induced pluripotent stem cells as a model system holds additional promise in investigating genes that are not as universally expressed across cell types as the genes we identify here, since they can be differentiated into a variety of specialized cell types.

## Conclusions

Here we identify a novel role for *ZNF335* in modifying circulating levels of LDL-cholesterol and LDL-cholesterol statin response by leveraging naturally occurring variation in ZNF335 expression in statin clinical trial participant-derived cell lines and a hypomorphic missense mouse model for *ZNF335*. Thus, it is possible that inter-individual variation in *ZNF335* expression contributes to differences in cholesterol levels and statin response in humans.

## List of Abbreviations

LDLC: low-density lipoprotein cholesterol
CVD: cardiovascular disease
GWAS: genome-wide association study
SREBF2: sterol regulatory element binding transcription factor 2
ZNF335: zinc finger protein 335 (human)
ZFP335: zinc finger protein 335 (murine)
LCL: lymphoblastoid cell line
CAP: Cholesterol and Pharmacogenetics
EBV: Epstein-Barr virus
AfrAm: African American
BMI: body mass index
RNA-seq: ribonucleic acid sequencing
ENU: N-ethyl-N-nitrosourea
TC: total cholesterol
HDLC: high-density lipoprotein cholesterol
IM: ion mobility
CCR4-NOT: carbon catabolite repression-negative on TATA-less
FDR: false discovery rate
ANOVA: analysis of variance
SEM: standard error of the mean
Q: false discovery rate adjusted p-value
F: female
M: male
IDL: intermediate density lipoprotein
VLDL: very low density lipoprotein

## Declarations

### Ethics approval and consent to participate

Written informed consent was obtained from all human participants, and the study was approved by the institutional review boards of the University of California, San Francisco and the University of California, Los Angeles. Mouse studies were approved by the Children’s Hospital & Research Center Oakland and the University of California, San Francisco Institutional Animal Care and Use Committees.

### Consent for publication

Not applicable.

### Availability of data and materials

CAP RNA-seq and phenotype data used for this work is available through the Cholesterol and Pharmacogenetics (CAP) Study dbGaP Study Accession: phs000481.v3.p2.

### Competing interests

The authors declare that they have no competing interests.

### Funding

This project was supported by NIH U19 HL069757, NIH P50 GM115318, AHA 15POST21880006, and AHA 12POST10430005. This study was also supported in part by the NIH Pharmacogenomics Research Network (PGRN) RNA Sequencing Project. Establishment and storage of the CAP LCLs was partially supported by NIH P30 DK063491. The funders had no role in the design of the study, the collection, analysis, or interpretation of the data, or in writing the manuscript.

### Authors’ contributions

ET helped to design the experiments, analyzed the data, and drafted the manuscript.

FYT conducted the mouse experiments, maintained the mouse line, and organized mouse data.

YQ assisted with mouse experiments.

KS and DN cultured LCLs and extracted RNA.

SMK supervised the mouse Liasys and ion mobility analysis.

NY assisted with mouse experiments.

JO performed mouse ion mobility analysis.

BYH and JGC generated the *Zfp335*^*R1092W*^ (bloto) mutant mouse line.

YIC and JIR contributed to the creation of the CAP LCLs.

RMK conducted the CAP clinical trial, helped interpret the ion mobility results, and revised the manuscript.

MWM supervised the work and revised the manuscript.

All authors read and approved the manuscript.

## Acknowledgements

We thank the CAP participants for their contributions to this study. We thank Arnie Acosta for culturing LCLs, Bahareh Sahami for lipid measurements, Jennifer Beckstead for mouse ion mobility measurements, Jinping An for conducting a pilot mouse study, Quest Diagnostics for providing the CAP IM data, and numerous UC Berkeley undergraduate volunteers for their assistance with this project. The mice used for this work were housed in a UCSF MLK Cores Research Facility. We fondly remember the late Dr. Debbie Nickerson and her contributions to this work while leading the Northwest Genomics Center.

## Supplementary Table Legends

**Table S1**. CAP LCL gene expression statin responses.

**Table S2**. Spearman correlations of statin-induced gene expression changes in LCLs with plasma LDL-cholesterol statin response in the corresponding CAP participants.

## Supplementary Figures

**Figure S1.**
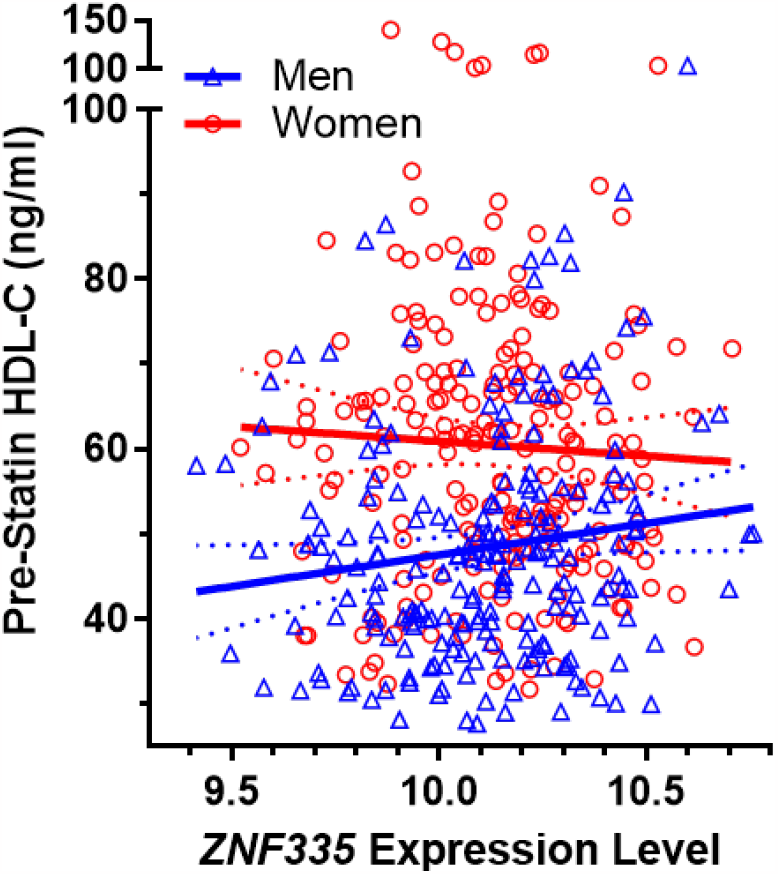
Correlation of pre-statin plasma HDL-cholesterol with *ZNF335* LCL gene expression levels from the corresponding donors split by sex (N=211 men, N=216 women).

**Figure S2.**
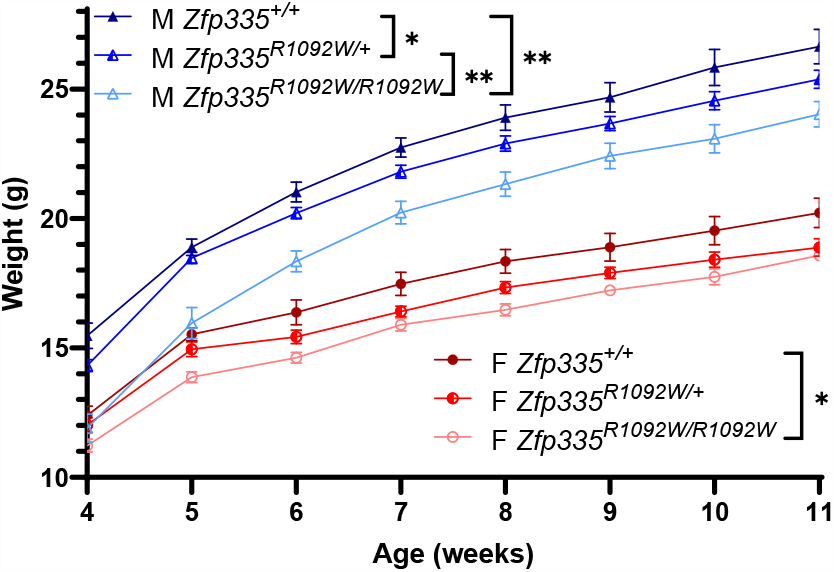
Mouse growth curves split by *Zfp335* genotype and sex. Adjusted p-values were calculated using the Compare Groups of Growth Curves (CGGC) method with 10,000 permutation tests. Male sample sizes were N=7, 19, and 7 and female sample sizes were N=7, 8, and 4 for wild type, heterozygotes, and homozygous Zfp335R1092W, respectively. Values are mean ± SEM. *p<0.05 **p<0.01

**Figure S3.**
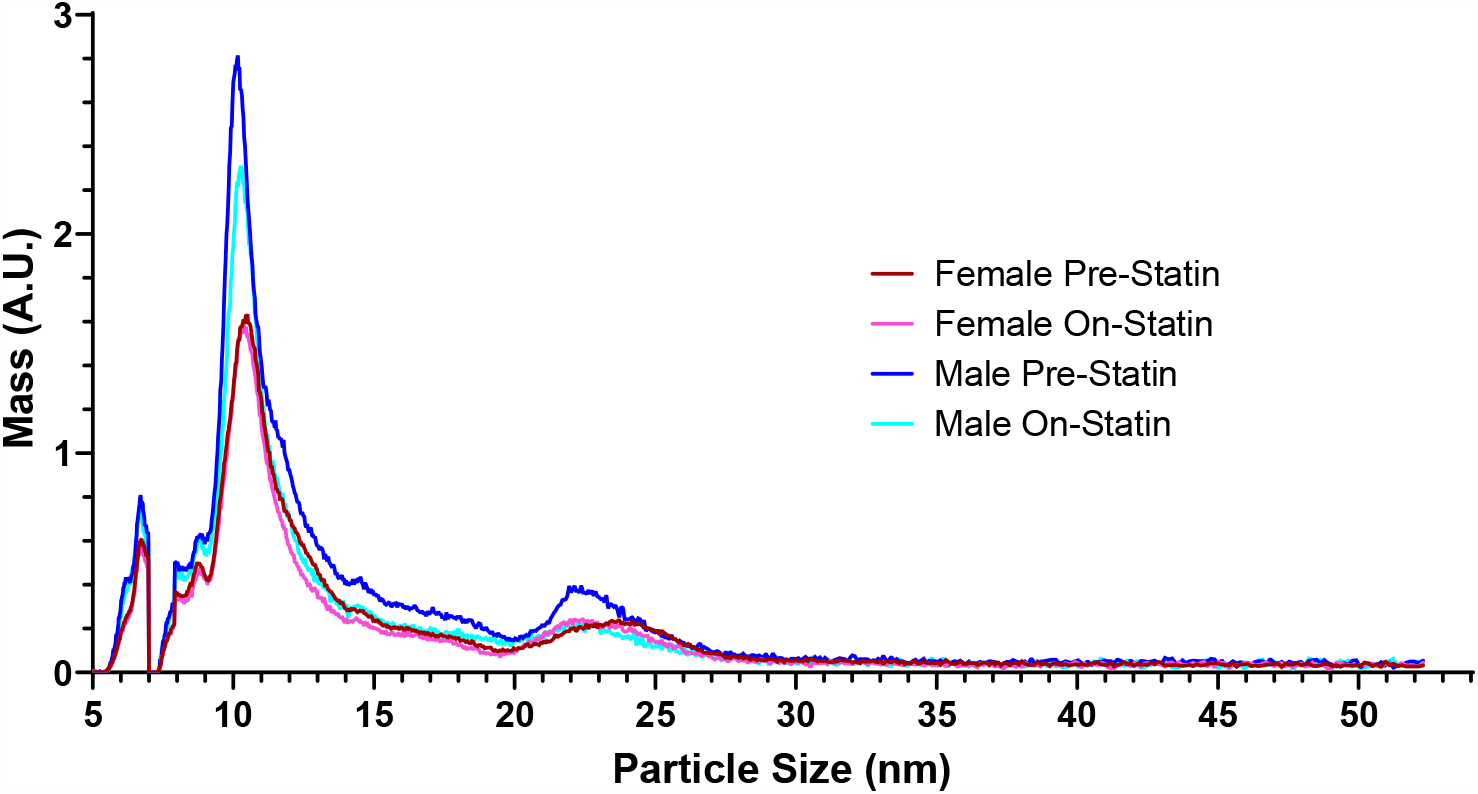
Statin-induced changes in wild-type N=5 male and N=11 female mouse lipoprotein profiles measured by ion mobility. Mouse profiles were measured before and after 4 weeks of simvastatin-containing diet. The size intervals designating the major lipoprotein subclasses are based on those defined in humans (29).

**Figure S4.**
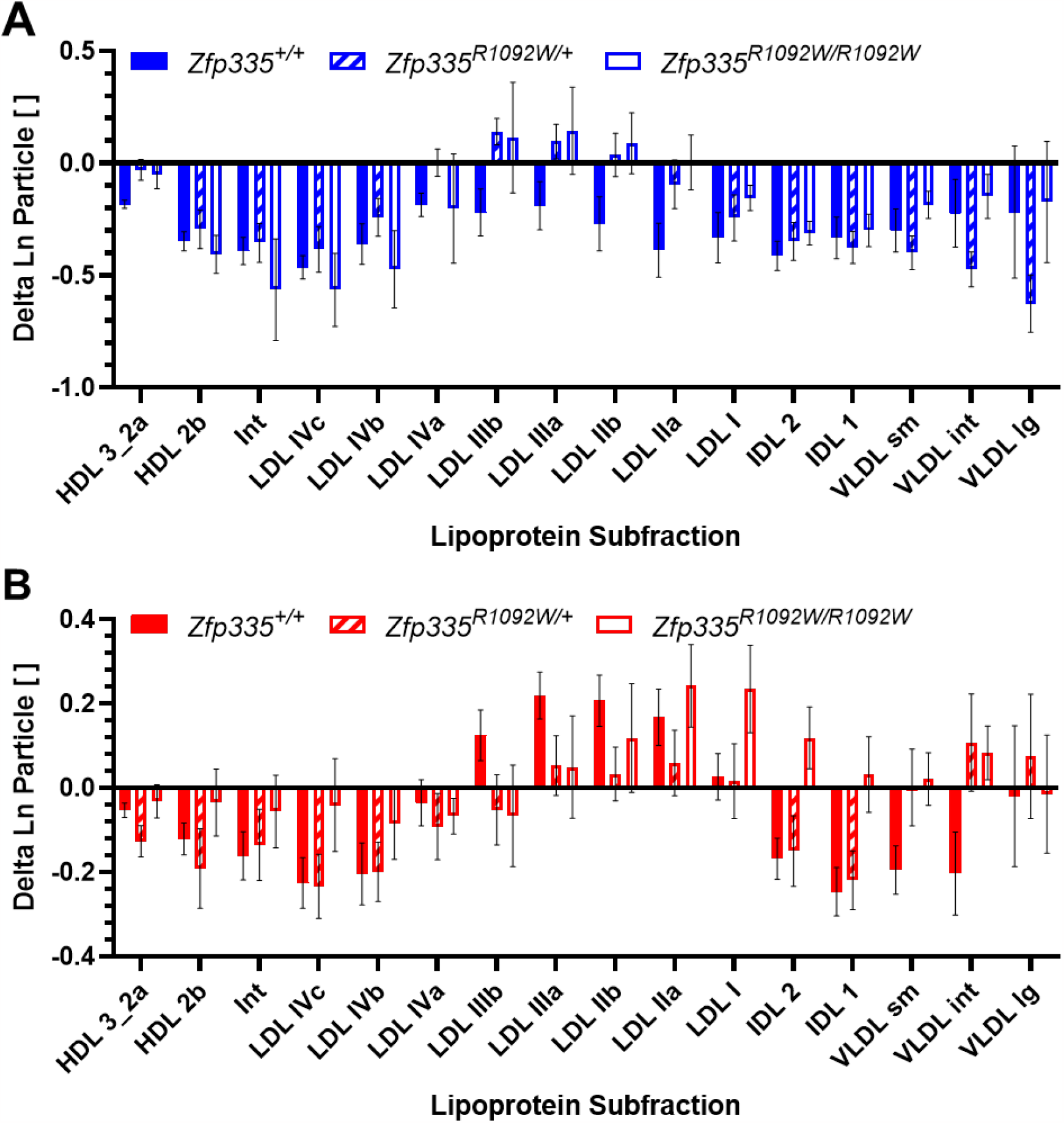
Statin-induced changes in (A) male and (B) female mouse plasma lipoprotein composition split by *Zfp335* genotype as measured by ion mobility. Male sample sizes were N=5, 12, and 5 and female sample sizes were N=11, 11, and 5 for wild type, heterozygotes, and homozygous Zfp335R1092W, respectively. Values are mean ± SEM.

**Figure S5.**
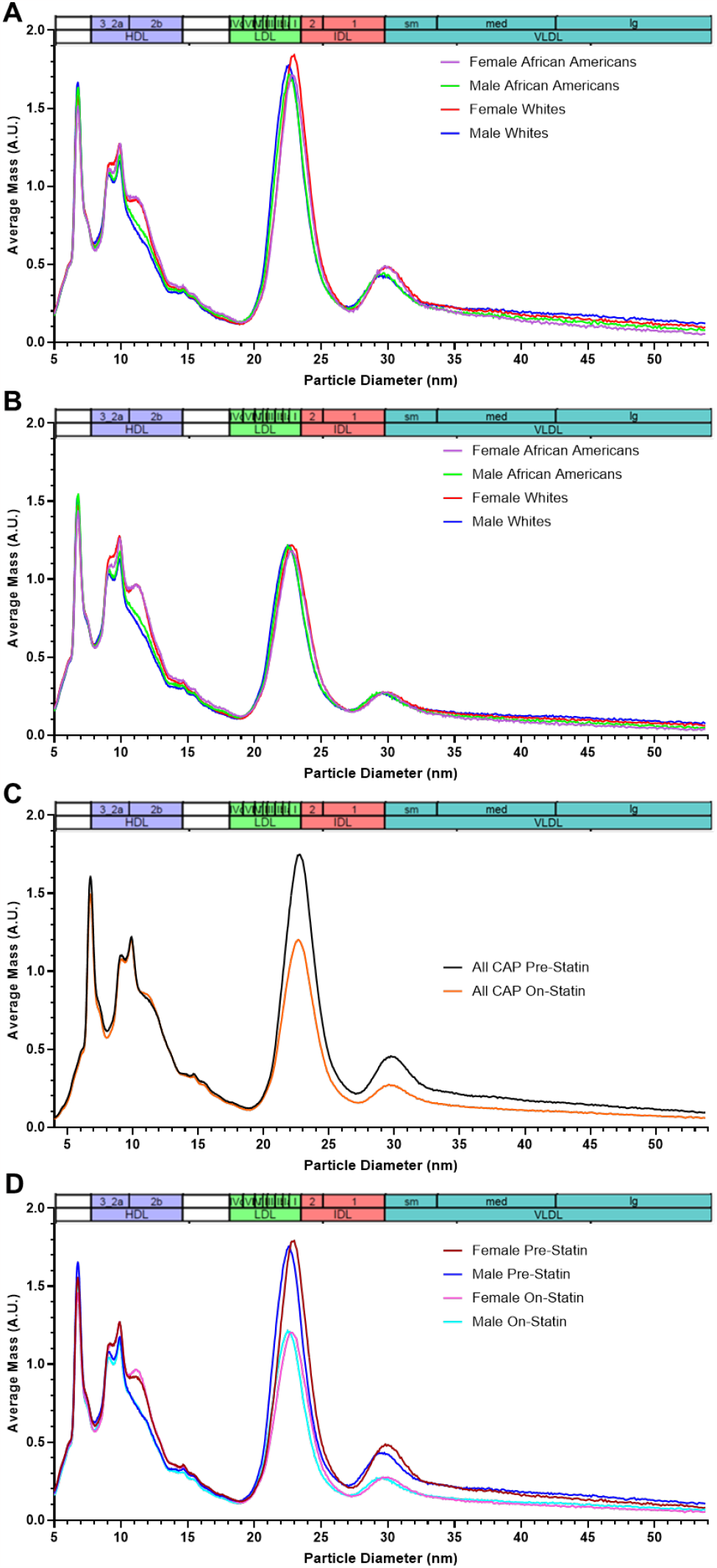
Ion mobility lipoprotein profiles for N=803 CAP participants before and after 6 weeks of 40 mg/day simvastatin treatment. Profiles shown are A) pre-statin split by sex and race/ethnicity B) on-statin split by sex and race/ethnicity C) pre- and on-statin for all and D) pre-statin and on-statin split by sex. N=142 for female African American, N=142 for male African American, N=243 for female white, and N=276 for male white participants.

**Figure S6.**
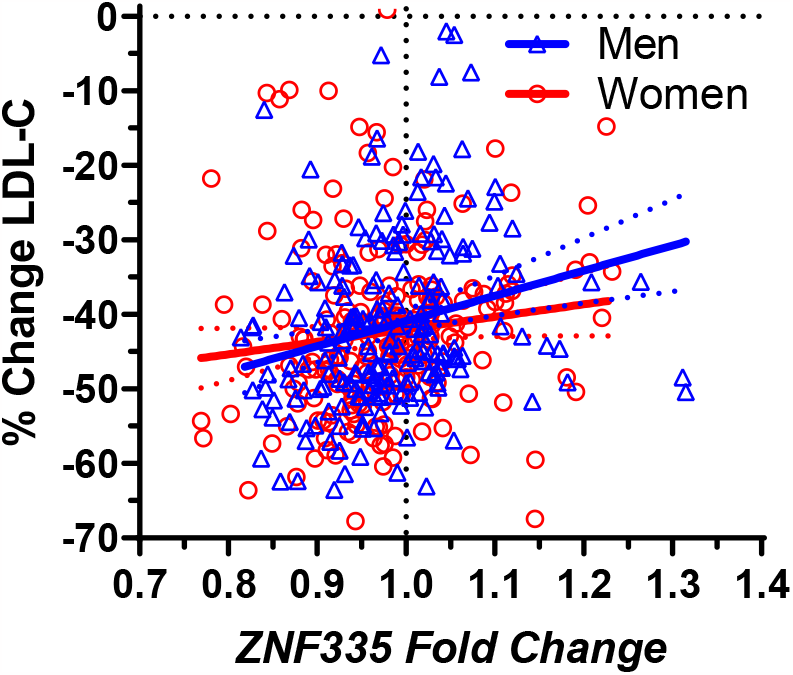
Correlation of plasma LDL-cholesterol statin response with *ZNF335* LCL gene expression statin response from the corresponding donors split by sex (N=211 men, N=216 women).

## Notes

### Competing Interest Statement

The authors have declared no competing interest.

## References

1. Arnett DK, Blumenthal RS, Albert MA, Buroker AB, Goldberger ZD, Hahn EJ, Himmelfarb CD, Khera A, Lloyd-Jones D, McEvoy JW, et al: 2019 ACC/AHA Guideline on the Primary Prevention of Cardiovascular Disease: A Report of the American College of Cardiology/American Heart Association Task Force on Clinical Practice Guidelines. Circulation 2019, 140:e596–e646.

2. Simon JA, Lin F, Hulley SB, Blanche PJ, Waters D, Shiboski S, Rotter JI, Nickerson DA, Yang H, Saad M, Krauss RM: Phenotypic predictors of response to simvastatin therapy among African-Americans and Caucasians: the Cholesterol and Pharmacogenetics (CAP) Study. Am J Cardiol 2006, 97:843–850.

3. Chasman DI, Giulianini F, Macfadyen J, Barratt BJ, Nyberg F, Ridker PM: Genetic Determinants of Statin-Induced Low-Density Lipoprotein Cholesterol Reduction: The Justification for the Use of Statins in Prevention: An Intervention Trial Evaluating Rosuvastatin (JUPITER) Trial. Circ Cardiovasc Genet 2012, 5:257–264.

4. Deshmukh HA, Colhoun HM, Johnson T, McKeigue PM, Betteridge DJ, Durrington PN, Fuller JH, Livingstone S, Charlton-Menys V, Neil A, et al: Genome-wide association study of genetic determinants of LDL-c response to atorvastatin therapy: importance of Lp(a). J Lipid Res 2012, 53:1000–1011.

5. Postmus I, Trompet S, Deshmukh HA, Barnes MR, Li X, Warren HR, Chasman DI, Zhou K, Arsenault BJ, Donnelly LA, et al: Pharmacogenetic meta-analysis of genome-wide association studies of LDL cholesterol response to statins. Nat Commun 2014, 5:5068.

6. Oni-Orisan A, Haldar T, Ranatunga DK, Medina MW, Schaefer C, Krauss RM, Iribarren C, Risch N, Hoffmann TJ: The impact of adjusting for baseline in pharmacogenomic genome-wide association studies of quantitative change. NPJ Genom Med 2020, 5:1.

7. Brown MS, Goldstein JL: The SREBP pathway: regulation of cholesterol metabolism by proteolysis of a membrane-bound transcription factor. Cell 1997, 89:331–340.

8. Medina MW, Theusch E, Naidoo D, Bauzon F, Stevens K, Mangravite LM, Kuang YL, Krauss RM: RHOA is a modulator of the cholesterol-lowering effects of statin. PLoS genetics 2012, 8:e1003058.

9. Mangravite LM, Engelhardt BE, Medina MW, Smith JD, Brown CD, Chasman DI, Mecham BH, Howie B, Shim H, Naidoo D, et al: A statin-dependent QTL for GATM expression is associated with statin-induced myopathy. Nature 2013, 502:377–380.

10. Kim K, Theusch E, Kuang YL, Dose A, Mitchel K, Cubitt C, Chen YI, Krauss RM, Medina MW: ZNF542P is a pseudogene associated with LDL response to simvastatin treatment. Sci Rep 2018, 8:12443.

11. Mitchel K, Theusch E, Cubitt C, Dosé AC, Stevens K, Naidoo D, Medina MW: RP1-13D10.2 Is a Novel Modulator of Statin-Induced Changes in Cholesterol. Circ Cardiovasc Genet 2016, 9:223–230.

12. Friedewald WT, Levy RI, Fredrickson DS: Estimation of the concentration of low-density lipoprotein cholesterol in plasma, without use of the preparative ultracentrifuge. Clin Chem 1972, 18:499–502.

13. Medina MW, Gao F, Ruan W, Rotter JI, Krauss RM: Alternative splicing of 3-hydroxy-3-methylglutaryl coenzyme A reductase is associated with plasma low-density lipoprotein cholesterol response to simvastatin. Circulation 2008, 118:355–362.

14. Theusch E, Kim K, Stevens K, Smith JD, Chen YI, Rotter JI, Nickerson DA, Medina MW: Statin-induced expression change of INSIG1 in lymphoblastoid cell lines correlates with plasma triglyceride statin response in a sex-specific manner. Pharmacogenomics J 2017, 17:222–229.

15. Parkhomchuk D, Borodina T, Amstislavskiy V, Banaru M, Hallen L, Krobitsch S, Lehrach H, Soldatov A: Transcriptome analysis by strand-specific sequencing of complementary DNA. Nucleic Acids Res 2009, 37:e123.

16. Theusch E, Chen YI, Rotter JI, Krauss RM, Medina MW: Genetic variants modulate gene expression statin response in human lymphoblastoid cell lines. BMC Genomics 2020, 21:555.

17. Kim D, Pertea G, Trapnell C, Pimentel H, Kelley R, Salzberg SL: TopHat2: accurate alignment of transcriptomes in the presence of insertions, deletions and gene fusions. Genome biology 2013, 14:R36.

18. Zerbino DR, Achuthan P, Akanni W, Amode MR, Barrell D, Bhai J, Billis K, Cummins C, Gall A, Giron CG, et al: Ensembl 2018. Nucleic Acids Res 2018, 46:D754–D761.

19. Arvey A, Tempera I, Tsai K, Chen HS, Tikhmyanova N, Klichinsky M, Leslie C, Lieberman PM: An atlas of the Epstein-Barr virus transcriptome and epigenome reveals host-virus regulatory interactions. Cell Host Microbe 2012, 12:233–245.

20. Anders S, Pyl PT, Huber W: HTSeq--a Python framework to work with highthroughput sequencing data. Bioinformatics 2015, 31:166–169.

21. Love MI, Huber W, Anders S: Moderated estimation of fold change and dispersion for RNA-Seq data with DESeq2. 2014.

22. Benjamini Y, Hochberg Y: Controlling the False Discovery Rate: A Practical and Powerful Approach to Multiple Testing. J Royal Stat Soc Ser B 1995, 57:289–300.

23. Han BY, Wu S, Foo CS, Horton RM, Jenne CN, Watson SR, Whittle B, Goodnow CC, Cyster JG: Zinc finger protein Zfp335 is required for the formation of the naive T cell compartment. Elife 2014, 3:e03549.

24. Elso CM, Roberts LJ, Smyth GK, Thomson RJ, Baldwin TM, Foote SJ, Handman E: Leishmaniasis host response loci (lmr1-3) modify disease severity through a Th1/Th2-independent pathway. Genes Immun 2004, 5:93–100.

25. Baldwin T, Sakthianandeswaren A, Curtis JM, Kumar B, Smyth GK, Foote SJ, Handman E: Wound healing response is a major contributor to the severity of cutaneous leishmaniasis in the ear model of infection. Parasite Immunol 2007, 29:501–513.

26. Chiu S, Bergeron N, Williams PT, Bray GA, Sutherland B, Krauss RM: Comparison of the DASH (Dietary Approaches to Stop Hypertension) diet and a higher-fat DASH diet on blood pressure and lipids and lipoproteins: a randomized controlled trial. Am J Clin Nutr 2016, 103:341–347.

27. Qin Y, Ting F, Kim MJ, Strelnikov J, Harmon J, Gao F, Dose A, Teng BB, Alipour MA, Yao Z, et al: Phosphatidylinositol-(4,5)-Bisphosphate Regulates Plasma Cholesterol Through LDL (Low-Density Lipoprotein) Receptor Lysosomal Degradation. Arterioscler Thromb Vasc Biol 2020, 40:1311–1324.

28. Musunuru K, Orho-Melander M, Caulfield MP, Li S, Salameh WA, Reitz RE, Berglund G, Hedblad B, Engström G, Williams PT, et al: Ion mobility analysis of lipoprotein subfractions identifies three independent axes of cardiovascular risk. Arterioscler Thromb Vasc Biol 2009, 29:1975–1980.

29. Caulfield MP, Li S, Lee G, Blanche PJ, Salameh WA, Benner WH, Reitz RE, Krauss RM: Direct determination of lipoprotein particle sizes and concentrations by ion mobility analysis. Clin Chem 2008, 54:1307–1316.

30. Garapaty S, Mahajan MA, Samuels HH: Components of the CCR4-NOT complex function as nuclear hormone receptor coactivators via association with the NRC-interacting Factor NIF-1. J Biol Chem 2008, 283:6806–6816.

31. Neely GG, Kuba K, Cammarato A, Isobe K, Amann S, Zhang L, Murata M, Elmén L, Gupta V, Arora S, et al: A global in vivo Drosophila RNAi screen identifies NOT3 as a conserved regulator of heart function. Cell 2010, 141:142–153.

32. Yang YJ, Baltus AE, Mathew RS, Murphy EA, Evrony GD, Gonzalez DM, Wang EP, Marshall-Walker CA, Barry BJ, Murn J, et al: Microcephaly gene links trithorax and REST/NRSF to control neural stem cell proliferation and differentiation. Cell 2012, 151:1097–1112.

33. Morita M, Oike Y, Nagashima T, Kadomatsu T, Tabata M, Suzuki T, Nakamura T, Yoshida N, Okada M, Yamamoto T: Obesity resistance and increased hepatic expression of catabolism-related mRNAs in Cnot3+/-mice. EMBO J 2011, 30:4678–4691.

34. Suzuki T, Kikuguchi C, Nishijima S, Nagashima T, Takahashi A, Okada M, Yamamoto T: Postnatal liver functional maturation requires Cnot complex-mediated decay of mRNAs encoding cell cycle and immature liver genes. Development 2019, 146.

35. Garapaty S, Xu CF, Trojer P, Mahajan MA, Neubert TA, Samuels HH: Identification and characterization of a novel nuclear protein complex involved in nuclear hormone receptor-mediated gene regulation. J Biol Chem 2009, 284:7542–7552.

36. Mahajan MA, Murray A, Samuels HH: NRC-interacting factor 1 is a novel cotransducer that interacts with and regulates the activity of the nuclear hormone receptor coactivator NRC. Mol Cell Biol 2002, 22:6883–6894.

37. The GTEx Consortium atlas of genetic regulatory effects across human tissues.Science 2020, 369:1318–1330.

38. Ratiu JJ, Barclay WE, Lin E, Wang Q, Wellford S, Mehta N, Harnois MJ, DiPalma D, Roy S, Contreras AV, et al: Loss of Zfp335 triggers cGAS/STING-dependent apoptosis of post-β selection thymocytes. Nat Commun 2022, 13:5901.

39. Liu H, Wang X, Ding R, Jiao A, Zheng H, Zhang C, Feng Z, Su Y, Yang X, Lei L, et al: The Transcription Factor Zfp335 Promotes Differentiation and Persistence of Memory CD8(+) T Cells by Regulating TCF-1. J Immunol 2022, 209:886–895.

40. Shirai YT, Suzuki T, Morita M, Takahashi A, Yamamoto T: Multifunctional roles of the mammalian CCR4-NOT complex in physiological phenomena. Front Genet 2014, 5:286.

41. Han BY, Foo CS, Wu S, Cyster JG: The C2H2-ZF transcription factor Zfp335 recognizes two consensus motifs using separate zinc finger arrays. Genes Dev 2016, 30:1509–1514.

